# Prevalence, Placenta Development, and Perinatal Outcomes of Women with Hypertensive Disorders of Pregnancy at Komfo Anokye Teaching Hospital

**DOI:** 10.1101/2020.05.14.095760

**Authors:** S. P. Awuah, I. Okai, E. A. Ntim, K. Bedu-Addo

## Abstract

**Background:** One of the most common medical problems associated with pregnancy is hypertension. Hypertensive disorders of pregnancy (HDP), which has been attributable to abnormal placentation may have adverse effects on both mother and foetus if left unchecked. The objective of this study was to determine the prevalence of Hypertensive Disorders of Pregnancy (HDP), the morphological variations of human placenta in HDP, and maternal and perinatal outcomes in HDP.

**Materials and Methods:** This was a prospective case-control study, conducted at Komfo Anokye Teaching Hospital (KATH), Ghana. The progression of pregnancy in normotensive and hypertensive pregnant women, and the eventual perinatal outcomes were closely followed. Statistical analysis was performed using IMB-SPSS version 23. Associations were considered significant at p values of ≤ 0.05.

**Results:** From a total of 214 deliveries recorded during the period of study, 84 (39.25%) were hypertensives. Forty four (52%) of the hypertensives had preeclampsia, 28 (33.3%) had gestational hypertension, 6 (7.1%) had eclampsia, 4 (4.8%) had chronic hypertension, and 2 (2.4%) had preeclampsia superimposed on chronic hypertension. The frequency of placental haematoma, placental infarction, and placental calcification in the normotensives were significantly (p=0.001) lower than that of the hypertensives. The mean placental weight (p = 0.01), placental volume (p = 0.001), placental diameter (p = 0.03), and placental thickness (p = 0.001) of the normotensives were significantly higher than those of the hypertensives. The number of normotensives in whom labour was induced, who had their babies delivered by caesarean section, and who were admitted after they had given birth were significantly (p=0.001) lower than that of hypertensives who underwent similar procedures. No stillbirths were recorded in the normotensives compared with four in the hypertensives. The number of babies delivered to the normotensives who were admitted to the NICU was significantly (p=0.001) lower than those delivered by hypertensives.

**Conclusion:** There was a high prevalence of hypertensive disorders of pregnancy in the study site. The condition adversely affected placental development and perinatal outcomes. These adverse effects can be curtailed by embarking on a vigorous health education drive.

## Introduction

Physiological changes occur in almost every pregnancy to aid in the nourishment and survival of the foetus. Biochemical parameters are good indicators of these adaptive changes in most organ systems and have shown to be different from the non-pregnant state [1]. These changes become very significant during complications of pregnancy. Hypertension is one of the medical problems that mostly affect pregnant women and it remains an important cause of both maternal and foetal morbidity/mortality. Studies show that 10-15% of pregnancies will be complicated by high blood pressure [2,3]. Up to about one-quarter of all antenatal admissions will be hypertensive related cases [2]. In developed countries, 16.1% of maternal deaths are as a result of hypertensive disorders and they are regarded as major risk factors for global maternal mortality [4]. Every year about 70,000 women die and there are half a million stillbirths or neonatal deaths owing to hypertensive disorders of pregnancy – the vast majority being in the developing world [5]. The identification of the disorder and its effective treatment play a beneficial role in pregnancy outcomes for the mother and the foetus, and hence a reduction in both maternal and perinatal mortality. Hypertensive disorder of pregnancy (HDP) has strong association with foetal growth retardation and preterm delivery, leading to perinatal morbidity or mortality. The placenta has been a valuable indicator for maternal and foetal diseases [6]. Many pregnancy complications which are associated with high foetal morbidity and mortality have shown gross deviations from the normal placenta morphology and anatomy [7]. Abnormal placenta adversely affects foetal outcome [8]. With the placenta serving as the image for the health status of the mother and foetus, complications like hypertension in pregnancy has reflected in the placenta in a significant way, either microscopically or macroscopically [9,10]. Pregnancies that are complicated by hypertension have been known to record higher incidence of neonatal morbidity compared to pregnancies with normal blood pressure. Pregnancies with hypertensive disorders are prone to a higher risk of preterm deliveries and low birth weights compared to healthy pregnancies [11]. Many investigators have reported that HDP has an adverse effect on newborn babies. The risk of HDP occurs mostly among mothers affected with severe chronic hypertension as well as those with superimposed preeclampsia on chronic hypertension [12]. The objective of this was to determine the prevalence of HDP, the morphological variations of human placenta in HDP, maternal and neonatal outcomes in HDP.

## Materials and methods

This was a prospective case-control study, conducted at the maternity block of Komfo Anokye Teaching Hospital (KATH) in Kumasi, Ghana, during the period of February 2018 to July 2018. Samples for the study were collected following approval from the Research and Development Unit, KATH and the Committee on Human Research, Publication and Ethics (CHRPE)-KNUST. Informed patient consent was obtained from all the participants after explaining the study in detail to them and ensuring confidentiality. Patients’ obstetric history were reviewed and those with hypertensive disorders of pregnancy were included in the study. Mothers with associated medical problems other than hypertension and those without antenatal records were excluded from the study.

The study participants were divided into four groups of hypertensive women according the classification system developed by the Working Group of the national high blood pressure education program [13]. These are *chronic hypertension, gestational hypertension, preeclampsia, eclampsia*, and *preeclampsia superimposed on chronic hypertension*.

Maternal parameters recorded/measured included the age of participants, body mass index, the final blood pressure reading before and after delivery, parity, occupation and level of income, educational level, ethnicity, other previous obstetric medical history, and booking status. Blood pressure of participants were recorded as a clinical routine using either a digital-portable automated blood pressure recorder or a standard mercury sphygmomanometer with the woman’s legs resting on a flat surface. The mode of delivery (vaginal or caesarean section), maternal presentation, and the degree of tear of maternal perineum were also recorded.

All the parameters of placenta were determined accurately using freshly delivered placentae from both the HDP group (hypertensive mothers) and the control group (normotensive mothers) at the labour ward, A1 HDU, and the theatre. Placentae were examined for haematomas, calcification or infarcts. The placental parameters measured included the weight, volume, diameter, and thickness, and shape. The umbilical cord insertion types, length, and the number of vessels in the umbilical cords were noted, along with any umbilical cord abnormalities.

Gestational age was expressed as beginning from the last date of menstruation proven by preliminary examination with ultrasound scan. On the basis of gestational age, the infants were categorized into 3 groups: *Term infants* were those with gestational age between 37 to 42 weeks, *preterm babies* included infants with gestational age <37 weeks and *post term babies* had gestational age >42 weeks. Low birth weight (LBW) was specified for birth weight (BW) <2.5 kg, very low birth weight (VLBW) as BW <1.5 kg, and extremely low birth weight (ELBW) as BW <1 kg [14]. The standard body length was defined as length of baby ranging from 46.9 to 54.9 cm [15]. The standard head circumference was defined as circumference of the head of baby ranging from 33 to 37 cm [16]. The standard abdominal circumference was defined as the circumference of the abdomen of baby ranging from 31 to 33 cm [14].

Data was analyzed using Statistical Package for Social Sciences (SPSS) version 23.0. Statistical analysis was performed with student t-test, and chi-square test. P value equal to or less than 0.05 was considered statistically significant.

## Results

Eighty four (39.25%) out of the 214 participants, had hypertensive disorders of pregnancy (HDP). Of the 84 deliveries with hypertensive disorders of pregnancy, 28 (33.3%) were gestational hypertension, 44 (52.4%) had preeclampsia, 6 (7.1%) had eclampsia, 4 (4.8%) had chronic hypertension, and 2 (2.4%) had preeclampsia superimposed on chronic hypertension (Fig. 1).

**Figure 1:**
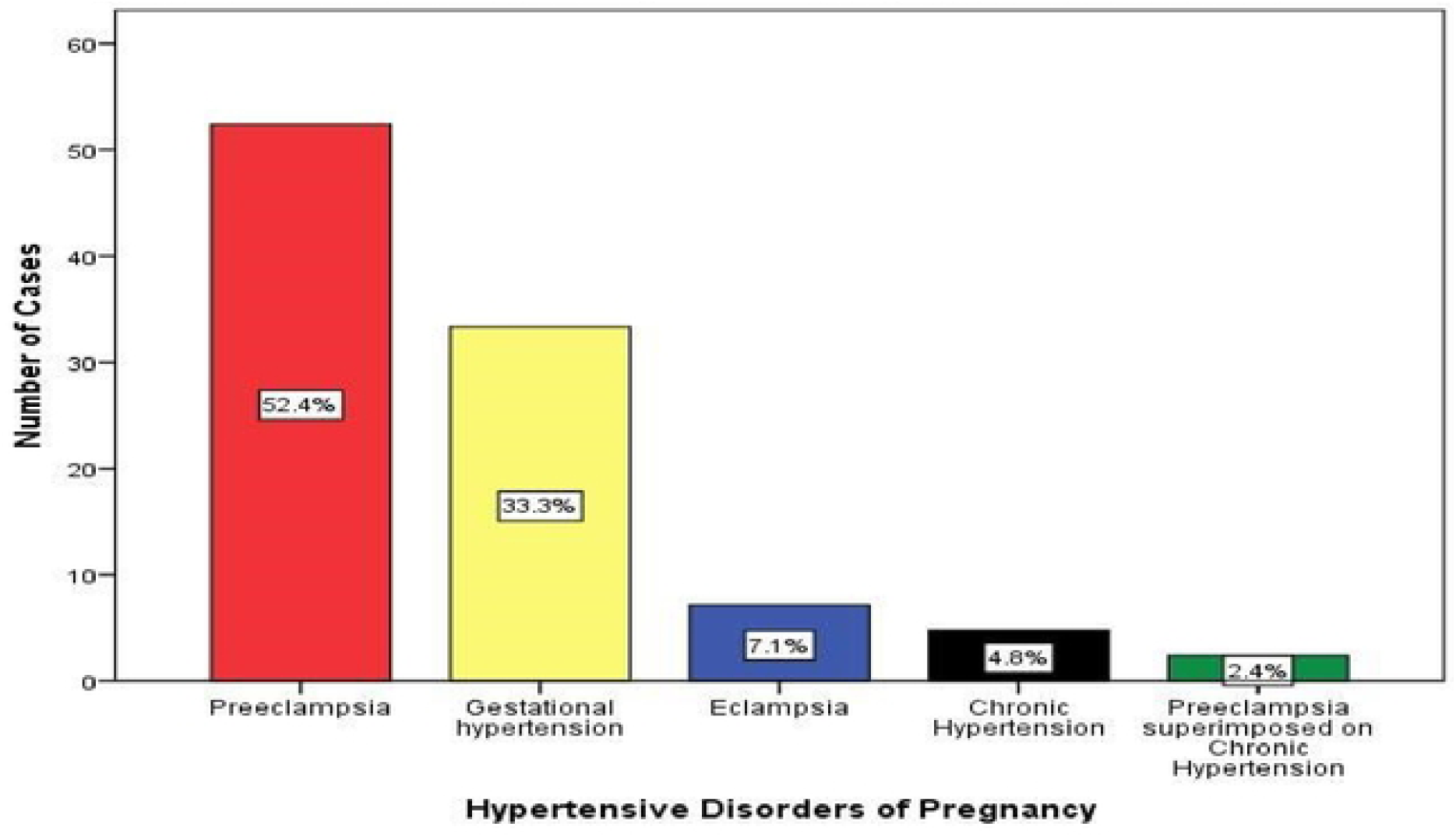
Distribution of cases HDP group according to type of hypertension.

The mean age of the hypertensives was 29.85 years. Among the hypertensives, 31 (36.9%) were in the age group 30-34 years, and 25 (29.8%) were in the age group 25-29 years. There was a significant difference in weight (p = 0.001) and height (p = 0.025) between the hypertensives and normotensives, but the difference in BMI observed between them was not significant (p = 0.090) (Table 1). The study showed that 57 (43.8%) normotensives and 42 (50.0%) of the hypertensives were obese.

**Table 1:** Distribution according to maternal age, Weight, Height, and BMI.

| Variable | Frequency |  | Mean±SD |  | ‘t’ | df | p-value |
| --- | --- | --- | --- | --- | --- | --- | --- |
| Age (years) | Contro<br>l | HDP | Control | HDP |  |  |  |
| ≤19 | 13 | 2 |  |  |  |  |  |
| 20-24 | 19 | 13 |  |  |  |  |  |
| 25-29 | 41 | 25 |  |  |  |  |  |
| 30-34 | 36 | 31 | 28.31±0.54 | 29.85±0.61 | 1.845 | 212 | 0.067 |
| 35-39 | 15 | 7 |  |  |  |  |  |
| ≥40 | 6 | 6 |  |  |  |  |  |
| MW (kg) | 130 | 84 | 72.79±13.10 | 78.84±12.46 | 3.368 | 212 | 0.001 |
| MH (m) | 130 | 84 | 1.58±0.08 | 1.60±0.07 | 2.255 | 212 | 0.025 |
| BMI (kg/m²) | 130 | 84 | 29.34±6.22 | 30.74±5.30 | 1.705 | 212 | 0.090 |

MW=Maternal Weight, MH=Maternal Height, BMI=Body Mass Index, SD=Standard Deviation, p=the p value.

Forty-eight (57.2%) of the hypertensives were multiparous, 19 (22.6%) primiparous and the remaining 17 (20.2%) were nulliparous. There was no significant difference in maternal parity between the normotensives and hypertensives (p = 0.324) (Table 2).

**Table 2:** Maternal parity.

| Parity | Frequency | | Percentage | | Sum of Frequencies | $\chi^2$ | df | p-value |
| --- | --- | --- | --- | --- | --- | --- | --- | --- |
|  | Control | HDP | Control | HDP |  |  |  |  |
| Para-0 | 39 | 17 | 30.0 | 20.2 | 56 | 3.476 | 3 | 0.324 |
| Para-1 | 32 | 19 | 24.6 | 22.6 | 51 |  |  |  |
| Para-2 | 29 | 22 | 22.3 | 26.2 | 51 |  |  |  |
| Para>3 | 30 | 26 | 23.1 | 31.0 | 56 |  |  |  |
| Total | 130 | 84 | 100.0 | 100.0 | 214 |  |  |  |

Majority of the hypertensives presented at the hospital (KATH) with headache as the chief complaint. During their hospitalization, the increases in diastolic blood pressure (DBP) corresponded with the systolic blood pressure (SBP) for both groups. The SBP was used as the major preliminary criteria for identifying the HDP group. There was a significant difference in the mean systolic BP (p = 0.001) and pulse rate (p = 0.001) of the normotensives and hypertensives before and after delivery (Table 3).

**Table 3:** Blood Pressure of Participants.

|  | SBP before delivery |  | SBP after delivery |  |
| --- | --- | --- | --- | --- |
|  | Control | HDP | Control | HDP |
| Minimum | 96 | 125 | 90 | 111 |
| Maximum | 128 | 240 | 110 | 185 |
| Mean±SD | 120.16±12.39 | 161.14±19.37 | 110.96±12.56 | 148.20±16.64 |
| 't' | 16.584 |  | 16.111 |  |
| p-value | 0.001 |  | 0.001 |  |

SBP=Systolic Blood Pressure, DBP=Diastolic Blood Pressure.

The number of normotensives in whom labour was induced, who had their babies delivered by caesarean section, and who were admitted after they had given birth were significantly (p=0.001) lower than that of hypertensives who underwent similar procedures (Table 4). No stillbirths were recorded in the normotensives compared with four in the hypertensives. The number of babies delivered by the normotensives who were admitted to the NICU was significantly (p=0.001) lower than those delivered by hypertensives. There was however, no significant differences in preterm deliveries, foetal presentation, and degree of maternal tear, between the two groups (Table 4).

**Table 4:** Delivery outcomes of study participants.

| Delivery outcome | Component | Control | | HDP | | '$\chi^2$' | df | p-value |
| --- | --- | --- | --- | --- | --- | --- | --- | --- |
|  |  | n | % | n | % |  |  |  |
| Induction of labour | Yes | 55 | 42.3 | 84 | 100.0 | 74.610 | 1 | 0.001 |
|  | No | 75 | 57.7 | 0 | 0.0 |  |  |  |
| Term of baby | Pre-term | 36 | 27.7 | 26 | 31.0 | 0.264 | 1 | 0.608 |
|  | Term | 94 | 72.3 | 58 | 69.0 |  |  |  |
| Mode of delivery | Vaginal | 128 | 98.5 | 37 | 44.0 | 85.581 | 1 | 0.001 |
|  | C/S | 2 | 1.5 | 47 | 56.0 |  |  |  |
| Presentation | Cephalic | 126 | 96.9 | 8 | 92.9 | 1.894 | 1 | 0.169 |
|  | Breech | 4 | 3.1 | 6 | 7.1 |  |  |  |
| Perineum of mothers | Intact | 74 | 57.4 | 47 | 56.0 | 0.020 | 1 | 0.886 |
|  | Tear | 55 | 42.6 | 37 | 44.0 |  |  |  |
| Type of birth | Live | 130 | 100.0 | 80 | 95.2 | 6.308 | 1 | 0.012 |
|  | IUFD | 0 | 0.0 | 4 | 4.8 |  |  |  |
| NICU admissions | Yes | 27 | 20.8 | 37 | 44.0 | 13.191 | 1 | 0.001 |
|  | No | 103 | 79.2 | 47 | 56.0 |  |  |  |
| Maternal admissions after delivery | Yes | 1 | 0.8 | 84 | 100.0 | 209.856 | 1 | 0.001 |
|  | No | 129 | 99.2 | 0 | 0.0 |  |  |  |

NICU=Neonatal Intensive Care Unit, C/S=Caesarean Section, IUFD=Intrauterine Foetal Death, p=the p value.

The frequency of placental haematoma, placental infarction, and placental calcification in the normotensives were significantly (p=0.001) lower than that of the hypertensives. One hundred and three (79.2%) of the placentae of the normotensives were oval shaped, compared to 30 (35.7%) of the hypertensives. Sixteen (12.3%) of the placentae of normotensives were oval compared to 41 (48.8%) for the hypertensives. The number of irregular shaped placentae was 11 (8.5%) and 13 (15.5%) for the normotensives and hypertensives respectively (Table 5).

**Table 5:** Gross morphology of Placenta of Study Participants.

| Parameter | Control |  | HDP |  | p-value |
| --- | --- | --- | --- | --- | --- |
|  | Frequency | % | Frequency | % |  |
| Haematoma |  |  |  |  |  |
| Yes | 5 | 3.8 | 28 | 33.3 | 0.001 |
| No | 125 | 96.2 | 56 | 66.7 |  |
| Infarction |  |  |  |  |  |
| Yes | 12 | 9.2 | 30 | 35.7 | 0.001 |
| No | 118 | 90.8 | 54 | 64.3 |  |
| Calcification |  |  |  |  |  |
| Yes | 43 | 33.1 | 51 | 60.7 | 0.001 |
| No | 87 | 66.9 | 33 | 39.3 |  |
| Placental Shape |  |  |  |  |  |
| Round | 103 | 79.2 | 30 | 35.7 | 0.01 |
| Oval | 16 | 12.3 | 41 | 48.8 |  |
| Irregular | 11 | 8.5 | 13 | 15.5 |  |

The mean placental weight (p = 0.01), placental volume (p = 0.001), placental diameter (p = 0.03), and placental thickness (p = 0.001) of normotensives were significantly higher than those of the hypertensives (Table 6).

**Table 6:** Placental Indices of the participants.

| Variable | Minimum | | Maximum | | Mean±SD | | 't' | 'df' | p-value |
| --- | --- | --- | --- | --- | --- | --- | --- | --- | --- |
|  | Control | HDP | Control | HDP | Control | HDP |  |  |  |
| **PW (kg)** | 0.20 | 0.20 | 0.74 | 0.60 | 0.56±0.13 | 0.49±0.18 | 3.518 | 212 | 0.01 |
| **PV (L)** | 0.22 | 0.18 | 0.77 | 0.65 | 0.54±0.11 | 0.46±0.15 | 4.577 | 212 | 0.001 |
| **PD (cm)** | 15.00 | 14.00 | 26.00 | 23.00 | 20.01±1.95 | 19.23±1.96 | 3.012 | 212 | 0.03 |
| **PT (cm)** | 1.10 | 0.90 | 2.70 | 2.30 | 2.09±0.29 | 1.88±0.33 | 4.834 | 212 | 0.001 |

PW=Placental Weight, PV=Placental Volume, PD=Placental Diameter, PT=Placental Thickness SD=Standard Deviation, p=the p value.

Eighty five (65.4%) of the placentae of the normotensives had eccentric cord insertion, 21 (16.2%) were central, 22 (16.9%) were marginal and 2 (1.5%) were velamentous. Thirty one (36.9%) placentae of the hypertensives had central insertion, 28 (33.3%) were eccentric, 23 (27.4 %) were marginal, and 2 (2.4%) were velamentous. Although majority of the umbilical cords from both groups had 3 vessels, the mean number of umbilical cords that had 3 vessels were significantly (p = 0.001) higher in the normotensives, compared to the hypertensives (Table 7). Majority (61.5%) of the umbilical cords of the normotensives had normal lengths whiles majority (53.6%) of the cords hypertensives were short. The mean cord length of the hypertensives (39.11±13.05 cm) was significantly lower than that of the normotensives (51.01±16.13 cm) (p = 0.001) (Table 7).

**Table 7:** Umbilical Cord Indices.

| Parameter | Control |  | HDP |  | p-value |
| --- | --- | --- | --- | --- | --- |
|  | Frequency | % | Frequency | % |  |
| Umbilical Cord Insertion |  |  |  |  |  |
| Velamentous | 2 | 1.5 | 2 | 2.4 | 0.001 |
| Central | 21 | 16.2 | 31 | 36.9 |  |
| Eccentric | 85 | 65.4 | 28 | 33.3 |  |
| Marginal | 22 | 16.9 | 23 | 27.4 |  |
| Umbilical Cord Vessel Number |  |  |  |  |  |
| Three | 121 | 93.1 | 59 | 70.2 | 0.001 |
| Two | 8 | 6.2 | 21 | 25.0 |  |
| One | 1 | 0.8 | 4 | 4.8 |  |
| Umbilical Cord Length (cm) |  |  |  |  |  |
| Short | 31 | 23.9 | 45 | 53.6 | 0.001 |
| Normal | 80 | 61.5 | 37 | 44.0 |  |
| Long | 19 | 14.6 | 2 | 2.4 |  |

The mean birth weights, birth lengths, head circumferences, and abdominal circumferences of the neonates of the normotensives were significantly higher than that of the neonates of the hypertensives (p = 0.001) (Table 8). Additionally, the Apgar score at the 5^th^ minute of birth of neonates of the normotensives was significantly higher than that of the neonates of the hypertensives (p = 0.001) (Table 8).

**Table 8:** Neonatal Indices of Study participants.

| Parameter | Minimum |  | Maximum |  | Mean±SD |  | ‘t’ | ‘df’ | p-value |
| --- | --- | --- | --- | --- | --- | --- | --- | --- | --- |
|  | Control | HDP | Control | HDP | Control | HDP |  |  |  |
| BW (kg) | 0.90 | 0.72 | 4.41 | 3.80 | 3.07±0.55 | 2.70±0.75 | 4.197 | 212 | 0.001 |
| BL (cm) | 38.0 | 33.0 | 57.0 | 53.0 | 48.27±4.46 | 45.18±4.42 | 4.976 | 212 | 0.001 |
| HC (cm) | 28.0 | 24.0 | 41.0 | 40.0 | 34.68±2.45 | 33.14±3.81 | 3.594 | 212 | 0.001 |
| AC (cm) | 37.0 | 20.0 | 40.0 | 21.0 | 32.60±3.11 | 30.76±3.90 | 3.814 | 212 | 0.001 |
| AS | 3/10 | 0/10 | 10/10 | 9/10 | 8.40±1.19 | 7.70±1.75 | 3.472 | 212 | 0.001 |

BW=Birth Weight, BL=Baby Length, HC=Head Circumference, AC=Abdominal Circumference, AS=Apgar Score, SD=Standard Deviation, p=the p value.

## Discussion

HDP has become a major health issue worldwide, and the prevalence varies from one country to another as well as in different institutions. This study showed a HDP prevalence of 39.25% at Komfo Anokye Teaching Hospital (KATH) during the study period. There has been a reported incidence of 1.5% to 22% of all pregnancies, which is dependent upon the population sampled and the definitions used [17,18,19,20,21]. The variation may be due to differences in genetic factors, socioeconomic status, racial differences, and some other demographic features such as maternal age and parity [19,20,21]. Another reason might be the differences in terminologies used in the study methodologies. Pregnancy Induced Hypertension (PIH) for instance, has led to a significant debate with misleading account in HDP prevalence, rendering the term PIH obsolete and no longer recommended in literature [17]. The prevalence of HDP with respect to age-group distribution was at its peak in women between 30-34 years, many of whom were diagnosed with either gestational hypertension or preeclampsia. The lowest number was recorded for mothers who were ≤19 years, followed by women who were ≥40 years (many of them being chronic hypertensives). Other studies found the highest proportion and lowest proportion of women with HDP between 25-29 years and ≥40 years [22], results inconsistent with that of the present study. Other researchers have reported an increased risk of HDP like preeclampsia in younger women who are 21 years or below [23,24].

Most of the women with hypertensive disorders of pregnancy in this study were multiparous (parity of 3 or more), followed by primiparous women. This is similar to some studies which found a higher prevalence of HDP among women with grand multiparity (5 or more) [22,23]. The mean Systolic BP before and after delivery of normotensives were significantly (p = 0.001) lower than that of the hypertensives, a result similar to findings of other studies [26,27]. Some investigators have found that women with HDP had a mid-trimester decrease, which was followed by a progressive rise in both systolic BP and diastolic BP between 30-45 days postpartum [28,29]. The factors influencing the development of high blood pressure may differ depending on the particular type of hypertensive disorder, the study population (ethnicity or race), family history of the individual [30], life style and eating habit of the individual [31], and most importantly the age and parity of the pregnant woman [32]. From the antenatal history, mothers who had higher BMI at the beginning of pregnancy or were overweight or obese during gestation, showed higher SBP and DBP values in all gestational trimesters until delivery. Contrary to the findings of this study, some previous studies have found underweight pregnant women to be at risk of hypertension development which result in delivery of preterm infants [33,34]. Indeed, pregnant women who are underweight or overweight are often at high risk, and therefore women who are thin or underweight are sometimes encouraged to put on weight before conception [35].

In this study, most of the placentae (48.8%) of the hypertensives were oval in shape, while most of the placentae (79.2%) of the normotensives were round in shape, a finding similar to that of other studies [36,37,10]. The outcome of the placental shapes in this study is different from that of other studies which found that the shape of placentae from both hypertensives and normotensives were either oval or round [38],^38^ or no significant difference (p > 0.05) in the number of different placental shapes of the normotensives and hypertensives [39]. The present study found a significantly high incidence of placental haematoma in hypertensives compared with the normotensives, a finding consistent with the findings of other studies [46,47]. A study has found an association between placental haematoma and low Apgar score and also an association between larger haematomas and IUFD, due in part to separation of a considerable part of the villi from the utero placental circulation [41]. The frequency of placental infarction between hypertensives and normotensives was significant (p = 0.001) in this study. This is consistent with the results obtained by other studies [36,42,43]. Placental infarcts are known to have an adverse effect on growth and development of the newborns [43]. The present study also observed a significantly high placental calcification in the hypertensives compared to the normotensives. This is similar to the findings of a study which concluded that the foetal outcome in terms of birth weight of newborns to mothers having PIH and calcification of placentae was poor when compared to the control group [44]. Another study found that the incidence of calcification was equal in the control and hypertensive groups [45]. It is noteworthy that calcification that is seen in the placenta shows an evidence of placental senescence or degeneration [46].

The mean placental weight, volume, thickness, and diameter for hypertensives were significantly lower than that of the control group in the present study (p < 0.05). Similar outcomes in placental weight have been reported by other studies [37,10,47,48,49,50]. This study showed that placental weight is a valuable parameter for predicting newborn weight, because a significant linear correlation was observed for both the hypertensives (r = 0.579, p = 0.001) and the normotensives (r = 0.630, p = 0.001). Similar relations have been shown by other researchers [51,7,52]. The present study recorded a significant (p < 0.001) reduction in the mean central thickness of placentae in the hypertensives compared to the normotensives. This finding is consistent with that of other studies [53,54,55,56,57,58]. The results for placental volume obtained in this study was similar to that obtained by other studies [59,60,61].

Majority of the umbilical cord lengths in the hypertensive mothers were significantly short compared to that of the normotensives in this study. Short umbilical cord lengths are associated with a high rate of foetal abnormalities, such as abdominal wall defects and defects in the extremities and spine [62]. They are also associated with unsatisfactory foetal state, central nervous system complications, and low Apgar and IQ scores [63]. A normal umbilical cord has two arteries and a vein and is covered by Wharton’s jelly. Changes may sometimes occur during pregnancy that result in abnormal number of umbilical cord vessels [64]. The number of umbilical cords with three vessels in the normotensives was significantly higher compared to the hypertensives in this study. Almost 30% of the umbilical cords of the hypertensives had less than 3 vessels compared to only 7% of the normotensives. The result of the present study is contrary to that of Saha *et al*. (2014) [65] who found 3 vessels in all their samples.

The umbilical cord insertion site to the placenta can be central, eccentric, marginal (battledore), or velamentous (membranous). More than 90% of term placentae insertions are central or eccentric. Marginal cord insertion (MCI) and velamentous cord insertions (VCI) are classified as abnormal placental cord insertions (PCI). VCI occurs in approximately 1% of singleton pregnancies and MCI in approximately 7% [62]. Central and eccentric cord insertions were the most common in both normotensives and hypertensives in the present study. Eighty one percent of the umbilical cord insertions in the normotensives in this study were either central or eccentric compared to 70.2% in the hypertensives. The frequency of marginal and velamentous cord insertions was higher in the hypertensives. This finding is similar but not to the same degree as that of other studies [66,67]. Abnormalities of the umbilical cord, related to morphology, placental insertion, number of vessels and primary tumors, can influence the perinatal outcome and may be associated with other fetal anomalies and aneuploidies [64].

The number of preterm deliveries for the normotensive and hypertensive mothers in the present study was not significantly different. A result contrary to that of Yadav *et al*. (1997) [68] who recorded significantly high numbers of preterm deliveries among the hypertensives compared to the normotensives. The need to induce labour or perform a caesarean section on the mothers was significantly higher (p = 0.001) in the hypertensives than the normotensives. The still birth rate was also significantly higher (p=0.012) in the hypertensives. These findings are similar to that of Yadav *et al*. (1997) [68]. The number of babies born to normotensives who needed NICU care was significantly lower compared to those of the hypertensives, results similar that of other studies [68,69,70]. It must be stated however that the frequencies in the present study were at times higher or lower than that of these studies.

The present study showed that the foetal development rate of the hypertensives were affected by adverse maternal and placental factors. The mean birth weight, baby length, abdominal circumference, and head circumference of neonates of the hypertensives were significantly (p = 0.001) lower compared to the normotensives in this study. Similar findings of LBW babies were observed in other studies [36,37,10,71]. The most significant determinant of foetal weight of the newborns from both the hypertensives and the normotensives in this study were the placental indices. The APGAR scores after five minutes of delivery was significantly (p≥0.001) higher in infants of the normotensives compared to the infants of the hypertensives. This is consistent with the findings of other studies [70,71].

## Conclusion

The study found a high prevalence of hypertensive disorders of pregnancy at the study site. This was associated with adverse placental growth and development as well as poor maternal and perinatal outcomes.

## Acknowledgements

The authors wish to appreciate the support of the Heads and nurses of various units of the Department of Obstetrics and Gynaecology of Komfo Anokye Teaching Hospital, Kumasi, Ashanti Region and all participants who voluntarily participated in the study.

## Supporting information

**Figure 1: Distribution of cases HDP group according to type of hypertension**

